# *CanDrivR-CS*: A Cancer-Specific Machine Learning Framework for Distinguishing Recurrent and Rare Variants

**DOI:** 10.1101/2024.09.19.613896

**Authors:** Amy Francis, Colin Campbell, Tom Gaunt

## Abstract

**Motivation:** Missense variants play a crucial role in cancer development, and distinguishing between those that frequently occur in cancer genomes and those that are rare may provide valuable insights into important functional mechanisms and consequences. Specifically, if common variants confer growth advantages, they may have undergone positive selection across different patients due to similar selection pressures. Moreover, studies have demonstrated the significance of rare mutations that arise as resistance mechanisms in response to drug treatment. This highlights the importance of understanding the role of both recurrent and rare variants in cancer. In addition to this, most existing tools for variant prediction focus on distinguishing variants found in normal and disease populations, often without considering the specific disease contexts in which these variants arise. Instead, they typically build predictors that generalise across all diseases. Here, we introduce *CanDrivR-CS*, a set of cancer-specific gradient boosting models designed to distinguish between rare and recurrent cancer variants.

**Results:** We curated missense variant data from the International Cancer Genome Consortium (ICGC). Cancer-type-specific models significantly outperformed a baseline pan-cancer model, achieving a maximum leave-one-group-out cross-validation (LOGO-CV) F1 score of up to 90% for *CanDrivR-SKCM (Skin Cutaneous Melanoma)* and 89% for *CanDrivR-SKCA (Skin Adenocarcinoma)*, compared to 79.2% for the baseline model. Notably, DNA shape properties consistently ranked among the top features for distinguishing recurrent and rare variants across all cancers. Specifically, recurrent missense variants frequently occurred in DNA bends and rolls, potentially implicating regions prone to DNA replication errors and acting as mutational hotspots.

**Availability and Implementation:** All training and test data, and Python code are available in our *CanDrivR-CS* GitHub repository: https://github.com/amyfrancis97/CanDrivR-CS.

## 1 Introduction

Recent advancements in next-generation sequencing have revolutionised the discovery of human genetic variants, revealing their prevalence across diverse genetic backgrounds in both health and disease contexts (30; 22; 2; 35). Despite the progress in identifying these variants, understanding their biological implications and contributions to disease phenotypes remains a significant challenge. Gaining further insights into their roles in disease is, therefore, crucial for advancing drug development and precision medicine.

In recent years, significant efforts have been made to develop methods for interpreting the role of single nucleotide variants (SNVs). Functional assays, such as Multiplexed Assays of Variant Effect (MAVE), are increasingly being used for this purpose (5; 14; 34; 20). MAVE assays systematically introduce genetic alterations and measure their impact on cellular phenotypes, including growth, gene expression, and protein function. However, the vast size of the human genome results in a combinatorial explosion, creating substantial challenges in terms of scalability, labour intensity, and cost. Consequently, there is an urgent need for complementary tools to assist in interpreting and predicting the effects of novel genetic variants within disease contexts.

In parallel, several computational tools have emerged to predict the pathogenic consequences of SNVs. Widely used tools such as *PolyPhen-2* (1), *SIFT* (19), and *MutationTaster* (28) assess the potential impact of SNVs across various diseases. However, these tools often lack specificity to variants that drive particular diseases, instead predicting pathogenic consequences across all disease genomes. Notably, our *CScape* predictor focused on predicting SNV impacts in cancers (27), yet without differentiation among cancer types.

In addition to understanding how mutational landscapes differ between cancer types, further insights could be gained by investigating why many cancer variants are rare while others occur more frequently. For example, rare secondary mutations can arise as mechanisms of resistance to treatment, such as the secondary resistance mutations that develop in the EGFR gene in response to Gefitinib treatment (21). Furthermore, common mutations may play a critical role in cancer development and progression, potentially undergoing positive selection across multiple individuals due to the growth advantage they confer to tumour cells; driving cancer phenotypes (16; 33).

This paper introduces *CanDrivR-CS*, a collection of cancer-specific machine learning algorithms that are designed to distinguish between recurrent and rare SNVs, trained on data from the International Cancer Genome Consortium (ICGC) (10). The methodology for classifying variants was inspired by our previous work in *CScape-Somatic* (26). Our findings suggest that building tailored machine learning models that consider the disease context of a variant can increase the performance by up to 11% (see Table 3 and Figure 1), compared to taking a pan-cancer prediction approach.

**Figure 1:**
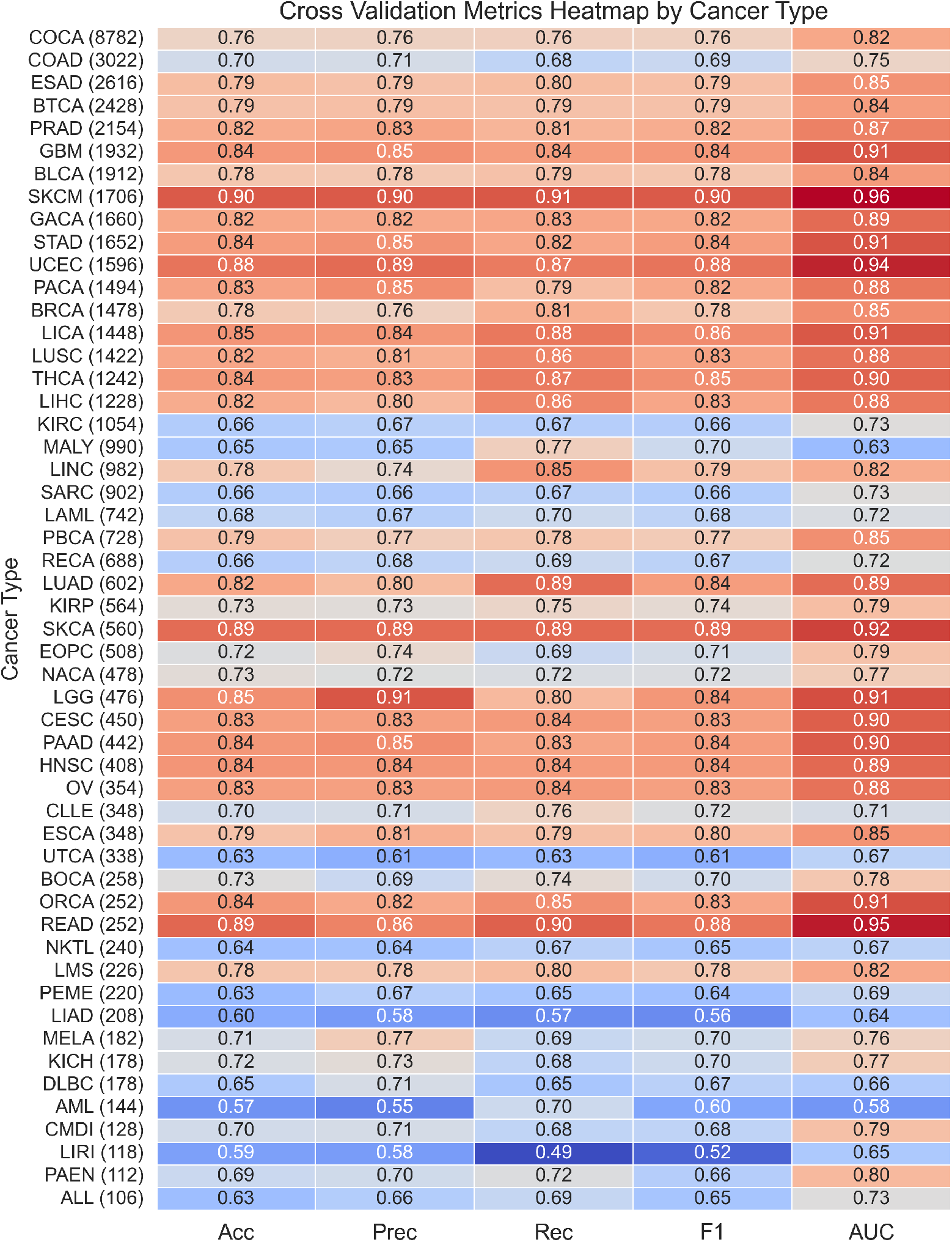
Cross-validation results heatmap sorted by total dataset size, with datasets below 100 samples excluded. Cancer datasets exceeding 1000 samples consistently lead to high predictive performance. Notably, UCEC, SKCM, and READ emerge as top performers, achieving F1 scores of 88%, 90%, and 88%, respectively. Despite their smaller sizes, datasets like SKCA also performed well.

## 2 Materials and Methods

The following section outlines the methods used in this work. For further descriptions, please refer to our GitHub repository.

### 2.1 Preparation of Train and Test Data

All datasets were prepared by filtering to autosomal, single nucleotide missense variants. Sex chromosomes and synonymous variants are functionally distinctive and would be the focus of a separate study. A summary of the training and test datasets used in this analysis are described in Table 1.

**Table 1:** This table shows a summary of the train and test datasets used in this analysis.

| Data | Variant Type | Data Type | No. Variants |
| --- | --- | --- | --- |
| ICGC | Somatic | Train/Test | 135,648 |
| COSMIC | Somatic | Test | 240,894 |
| TCGA-UCEC | Somatic | Test | 557 |
| TCGA-SKCM | Somatic | Test | 103 |

#### 2.1.1 Training Data: The International Cancer Genome Consortium (ICGC)

##### Downloading ICGC Data

We downloaded V1.0 of the simple somatic mutation (aggregated) VCF file from the International Cancer Genome Consortium (ICGC) website (10). We updated the genomic coordinates from the hg19 genome build to the hg38 genome build using pyliftover in Python, facilitated by the hg19ToHg38 chain file from the UCSC Genome Browser (18).

##### Preparation of ICGC Data

In this work, we propose two models. The first model, *CanDrivR*, serves as our baseline model. This model takes a pan-cancer approach; designed to distinguish between recurrent and rare somatic mutations across all cancer types included in the ICGC dataset.

Building upon the baseline, we introduce *CanDrivR-CS*, an ensemble of cancer-specific machine learning models tailored to each individual cancer type within ICGC.

##### CanDrivR Baseline Model

For our baseline model, our rare class was defined as variants occurring in exactly one patient in the ICGC dataset. To establish the recurrent class, we experimented with training our model using different donor count thresholds (Supplementary Figure 1). *CanDrivR’s* performance was highest when using a donor count threshold of *>* 2 for the recurrent class. After applying this threshold, we identified 67,824 variants in the recurrent class and 2,035,826 variants in the rare class. To balance the data, we randomly down-sampled the rare class to match the number of recurrent variants. Hence, our final dataset contained 135,648 missense variants.

##### CanDrivR-CS Model

After developing our baseline model, we developed *CanDrivR-CS* by training a separate model for each unique cancer type within ICGC. To keep the sizes of our datasets large enough to model, we pooled cancer datasets across different study sub-populations. In *CanDrivR-CS* models, the rare class was defined in the same way as our baseline model; using variants with a donor count of exactly one. For the recurrent class, we optimised the donor count threshold for each cancer model. To determine these thresholds, we produced optimisation curves (see Supplementary Figure 2 for some examples) similar to those in the baseline model (Supplementary Figure 1). This approach allowed us to tailor each model to the specific mutation recurrence patterns observed in each type of cancer. We then down-sampled the majority class and removed any cancer types that had a balanced dataset of less than 100 samples, leading to 50 different cancer models. The thresholds and total dataset sizes used for training each of the *CanDrivR-CS* models are detailed in Table 2.

**Table 2:** Cancer Types, Donor Counts, and Sizes of Balanced Datasets. This table details the cancer codes used to build models based on the ICGC dataset, the thresholds selected for the recurrent training dataset in *CanDrivR-CS*, and the final column shows the total size of the balanced dataset used for training and testing. The definitions of cancer codes are derived from several sources (13; 18; 9; 10).

| Cancer Name | Cancer Code | Donor Count Threshold (>) | Size |
| --- | --- | --- | --- |
| Acute Lymphoblastic Leukemia | ALL | 1 | 106 |
| Acute Myeloid Leukemia | AML | 3 | 144 |
| Bladder Urothelial Carcinoma | BLCA | 2 | 1912 |
| Bone Cancer | BOCA | 2 | 258 |
| Breast Cancer | BRCA | 3 | 1478 |
| Biliary tract cancer | BTCA | 2 | 2428 |
| Cervical Squamous Cell Carcinoma | CESC | 3 | 450 |
| Chronic Lymphocytic Leukemia | CLLE | 2 | 348 |
| Colon Adenocarcinoma | COAD | 1 | 3022 |
| Colorectal Cancer | COCA | 6 | 8782 |
| Diffuse Large B-Cell Lymphoma | DLBC | 2 | 178 |
| Early Onset Prostate Cancer | EOPC | 2 | 508 |
| Esophageal Adenocarcinoma | ESAD | 3 | 2616 |
| Esophageal Carcinoma | ESCA | 4 | 348 |
| Gastric cancer | GACA | 3 | 1660 |
| Glioblastoma Multiforme | GBM | 3 | 1932 |
| Head and Neck Squamous Cell Carcinoma | HNSC | 2 | 408 |
| Kidney Chromophobe | KICH | 1 | 178 |
| Kidney Renal Clear Cell Carcinoma | KIRC | 1 | 1054 |
| Kidney Renal Papillary Cell Carcinoma | KIRP | 2 | 564 |
| Liver Adenocarcinoma | LIAD | 1 | 208 |
| Liver Intrahepatic Cholangiocarcinoma | LICA | 3 | 1448 |
| Liver Hepatocellular Cancer | LIHC | 2 | 1228 |
| Liver Cancer | LIRI | 1 | 118 |
| Lower Grade Glioma | LGG | 3 | 476 |
| Lung Adenocarcinoma | LUAD | 3 | 602 |
| Liver Cancer | LINC | 2 | 982 |
| Lung Squamous Cell Carcinoma | LUSC | 3 | 1422 |
| Leiomyosarcoma | LMS | 2 | 226 |
| Malignant Lymphoma | MALY | 1 | 990 |
| Melanoma | MELA | 2 | 182 |
| Nasopharyngeal Cancer | NACA | 1 | 478 |
| Natural Killer/T-cell Lymphoma | NKTL | 1 | 240 |
| Oral Cancer | ORCA | 2 | 252 |
| Ovarian Cancer | OV | 3 | 354 |
| Pancreatic Adenocarcinoma | PAAD | 3 | 442 |
| Pancreatic Cancer Endocrine neoplasms | PAEN | 2 | 112 |
| Pancreatic Cancer | PACA | 2 | 1494 |
| Pediatric Brain Cancer | PBCA | 2 | 728 |
| Pediatric Medulloblastoma | PEME | 2 | 220 |
| Prostate Cancer | PRAD | 2 | 2154 |
| Rectum Adenocarcinoma | READ | 6 | 252 |
| Rectal Cancer | RECA | 1 | 688 |
| Sarcoma | SARC | 1 | 902 |
| Skin Adenocarcinoma | SKCA | 5 | 560 |
| Skin Cutaneous Melanoma | SKCM | 4 | 1706 |
| Stomach Adenocarcinoma | STAD | 3 | 1652 |
| Thyroid Carcinoma | THCA | 4 | 1242 |
| Uterine Carcinosarcoma | UTCA | 1 | 338 |
| Uterine Corpus Endometrial Carcinoma | UCEC | 3 | 1596 |

#### 2.1.2 Baseline Model Test Data: The Catalogue of Somatic Mutations in Cancer (COSMIC)

We evaluated our *CanDrivR* baseline model on rare and recurrent mutations from the Catalogue of Somatic Mutations in Cancer (COSMIC) (32). Specifically, we downloaded v.99 of the COSMIC ‘genome screens mutant’ file in GRCh38 format. We then removed any variants present in our ICGC training data, filter to r = 1 and r *>*2, and down-sample the majority group. We remove all variants present in our training data. In total, this led to a dataset size of 240,894 variants.

#### 2.1.3 Cancer-Specific Test Data: The Cancer Genome Atlas (TCGA)

To evaluate our top-performing cancer-specific models, we used data from The Cancer Genome Atlas (TCGA) (9), selecting models with sufficient samples (over 100 after removing variants present in our training data). TCGA was chosen for its consistent cancer labelling with ICGC, enabling straightforward subgrouping by cancer type; minimising any bias. Using the *TCGAbiolinks* package (7) in R, we downloaded TCGA data.

We created unique mutation identifiers, and counted distinct tumour sample barcodes for each mutation. We focused on uterine corpus endometrial carcinoma (UCEC) and skin cutaneous melanoma (SKCM), as these were the only cancers with over 100 variants after excluding ICGC training data. To create balanced datasets, we filtered mutations by sample count thresholds (*>*4 for SKCM and *>*3 for UCEC) and randomly selected an equal number of mutations with a sample count of one, matching the thresholds in the respective *CanDrivR-CS* training data. This resulted in final dataset sizes of 103 variants for SKCM and 557 variants for UCEC.

### 2.2 Algorithms and Hyperparameters

For modeling, we used the `XGBClassifier()` function from the XGBoost library (4). To ensure reproducibility, we set the random state to 42 across all models and used the default hyperparameters for the `XGBClassifier` in all training processes. We opted for the default hyperparameters due to the extensive number of models (50 cancer-specific models plus a baseline model) that needed to be trained. Optimising hyperparameters for each model would have significantly increased computational complexity and time.

### 2.3 Handling Missing Data

XGBoost handles missing data using a sparsity-aware algorithm that learns the optimal direction for missing values during tree building process. Hence, we did not impute missing data ourselves (4). When encountering a missing value, XGBoost evaluates both possible split directions and selects the one that maximises gain (4).

### 2.4 Features

We extracted features using the *DrivR-Base* framework (11). For an in-depth description of the features and their sources, please refer to our supplementary material, and our previous *DrivR-Base* paper (11). In this work, we used sequential feature selection to retain the most informative features (see Supplementary Figure 3 for more details). Our feature selection led to a final feature set of 351 features. A full list of selected features can be found in Supplementary Tables 1-6.

The selected features encompass a wide range of biological properties, and can be broadly categorised into the following feature groups:

1. **Conservation-based scores**: These scores measure the evolutionary conservation of nucleotide sequences, indicating the likelihood of a variant being deleterious if it occurs in a highly conserved region (25; 29).
2. **Variant Effect Predictor consequences and amino acid prediction**: This group includes the predicted impact of variants, such as the introduction of stop codons (17).
3. **Dinucleotide properties**: Features related to the physical and chemical properties of dinucleotide sequences, which can influence DNA stability (12).
4. **DNA shape properties**: Structural properties of DNA, such as minor groove width and helix twist, which can affect how DNA interacts with proteins and other molecules (6).
5. **GC/CpG content**: The proportion of guanine-cytosine pairs and the presence of CpG islands, which are regions rich in CG dinucleotides and can be associated with gene regulatory elements.
6. **Kernel-based sequence similarity**: Computational measures that measure the similarity between sequences using kernel-based methods (3).
7. **Amino acid substitution matrices**: Matrices that provide scores for the likelihood of one amino acid being substituted for another, based on evolutionary data (e.g., BLOSUM, PAM) (24).
8. **Amino acid properties**: Features that describe the physical, chemical, and functional properties of amino acids, such as hydrophobicity, charge, and molecular weight (15).

#### 2.5 Model Evaluation

All models were optimised and evaluated using leave-one-group-out cross-validation (the definition of a *group* is given below). The rationale behind this choice was to prevent data leakage during the evaluation process. If we were to randomly partition samples into training and validation sets, it might result in training and test variants that are in close proximity and therefore share very similar features. This would not provide a true reflection of the model’s ability to generalise to unseen data. To mitigate this issue, we divided the data into 11 groups, with each group containing variants from two randomly assigned chromosomes. This ensured that our validation and test data remained distinct from our training dataset:

1. We randomly split our data into 11 groups, each containing variants from two distinct chromosomes (e.g., chr1 & chr5 variants assigned to group 1).
2. Out of these 11 groups, we then randomly selected a single group to hold out for testing.
3. From the remaining training/validation dataset, we held out a second group for validation. We trained our model using the remaining nine groups and validated our model using this held-out validation dataset.
4. We calculated the accuracy, precision, recall, F1, and area under the curve (AUC) scores, using XGBoost’s built in ‘metrics’ function.
5. We repeated this cross-validation process ten times, each with a different validation dataset.
6. Our final *validation* result is the mean of all metrics over all folds of cross-validation.
7. We then re-trained our model using all ten training/validation groups and tested on the 11th group. These results are defined as our *test* results.

## 3 Results

### 3.1 Baseline Model

#### 3.1.1 Evaluating on ICGC Data

We built our baseline gradient boosting model, *CanDrivR*, using pan-cancer variants from ICGC (10). As detailed in our Methods section, we evaluated our model using leave-one-group-out cross-validation (LOGO-CV). The mean metrics over all folds of cross-validation, along with the overall test performance, are presented in Table 3. *CanDrivR* was able to generalise very well across all chromosomes (crossvalidation F1 = 79.2 ± 1.7%) and the held out test set (F1 = 79.6%).

**Table 3:** Cross-validation and test results for rare (donor count = 1) and recurrent (donor count *>*2) variants.The cross-validation metrics represent the mean results for the cross-validation folds, and the standard deviations represent the standard deviations across the folds. The test statistics represent the held-out test data, and unseen COSMIC data.

| Dataset | Result | Acc. (%) | Prec. (%) | Rec. (%) | F1 (%) | AUC (%) |
| --- | --- | --- | --- | --- | --- | --- |
| ICGC | Cross-Val | $78.9 \pm 1.5$ | $78.4 \pm 1.7$ | $80.0 \pm 2.3$ | $79.2 \pm 1.7$ | $85.2 \pm 1.8$ |
| ICGC | Test | 80.1 | 78.3 | 80.8 | 79.6 | 79.8 |
| COSMIC | Test | 66.5 | 64.5 | 66.7 | 65.6 | 65.0 |

#### 3.1.2 Evaluating on ‘Unseen’ COSMIC Data

Next, we evaluated our baseline model using data from COSMIC (32). Our baseline model was unable to generalise well to this unseen dataset, and only achieved an F1 score of 65.6% (Table 3). We discuss possible explanations for this observation in the discussion.

### 3.2 Cancer-Specific Models

#### 3.2.1 Evaluation

Following our baseline model, we developed *CanDrivR-CS*, a cancer-specific gradient boosting framework. We partitioned the ICGC dataset into smaller subsets based on cancer type and constructed distinct models for each of these. *CanDrivR-CS* consistently achieved markedly higher cross-validation performance, with the top models reaching an F1 score of 90% (Figure 1), compared to the baseline model’s F1 score of 79.2%. Notably, *CanDrivR-SKCM (Skin Cutaneous Melanoma)* and *CanDrivR-SKCA (Skin Adenocarcinoma)* were among the highest performers, achieving F1 cross-validation scores of 90% and 89%, respectively.

Furthermore, datasets exceeding 1000 samples consistently achieved the highest performance, with most F1 scores surpassing 79%. In contrast, smaller datasets containing fewer than 1000 variants showed greater variability. For instance, *CanDrivR-LIRI (Liver Cancer)* yielded an F1 score of 52%, with a dataset size of 118 variants. However, there were exceptions to this trend; *CanDrivR-COAD (Colorectal Cancer)* only achieved an F1 score of 69%, despite its dataset size exceeding 3000 variants. Overall, our results underscore a significant gain in prediction accuracy when considering the cancer-specific context in which variants occur.

#### 3.2.2 Feature Importance

We next investigated the most important features for distinguishing rare and recurrent variants. We compared the top five features for the top-performing cancer-specific models (leading to a comparison of 32 features across ten cancers). We present the results as a heatmap in Figure 2.

**Figure 2:**
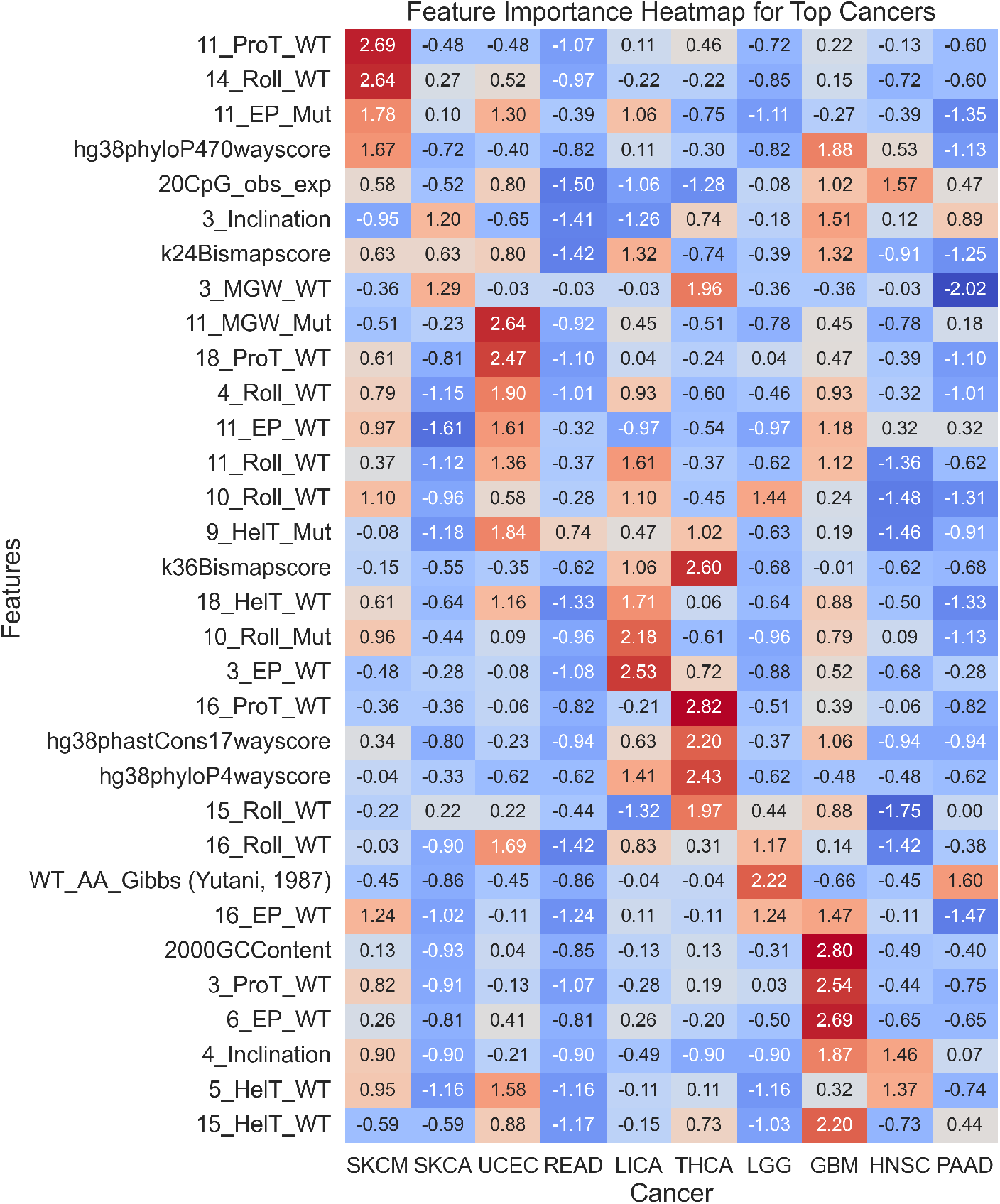
Here, we calculated the feature importance for the top five features for each of the cancer types. The feature importance was calculated using XGBoost’s get booster().get score() function, and then the data was scaled using Scikit Learn’s StandardScaler() for plotting. For each of the DNA shape features (E.g., Propellar Twist (ProT), Helix Twist (HelT), Electrostatic Potential (EP), and Roll), the number proceeding the feature type is the position at which the value is recorded. For example, a position of ‘11’ indicates the site of the nucleotide variant, a position of ‘10’ indicates the nucleotide position to the left of the variant, and ‘12’ indicates the nucleotide immediately to the right of the variant. For more information on DNA shape features, please see Supplementary Tables 1-6.

The top features included DNA shape properties and conservation scores (6). Despite the helix twist (HelT), propeller twist (ProT), electrostatic potential (EP), and minor groove width (MGW) DNA shape features being informative for most cancer types, their relative importance varied by sequence position. For instance, in Skin Cutaneous Melanoma (SKCM), the ProT value is most important at the site of substitution in the wild-type sequence (denoted by the value ‘11’), whereas, for UCEC, the importance of ProT is more pronounced at position 18 in the mutant sequence. This variability explains why cancer-specific models are unable to generalise well to other cancer datasets (see Supplementary Figure 6).

We investigated DNA shape features further for SKCA and SKCM datasets, by plotting the mean values at each position, in both wild type and mutant sequences, for both recurrent and rare variants (Figure 3). We found that recurrent variants occurred in DNA regions that were more bent and twisted. Conversely, rare variants exist within ‘flatter’ DNA regions.

**Figure 3:**
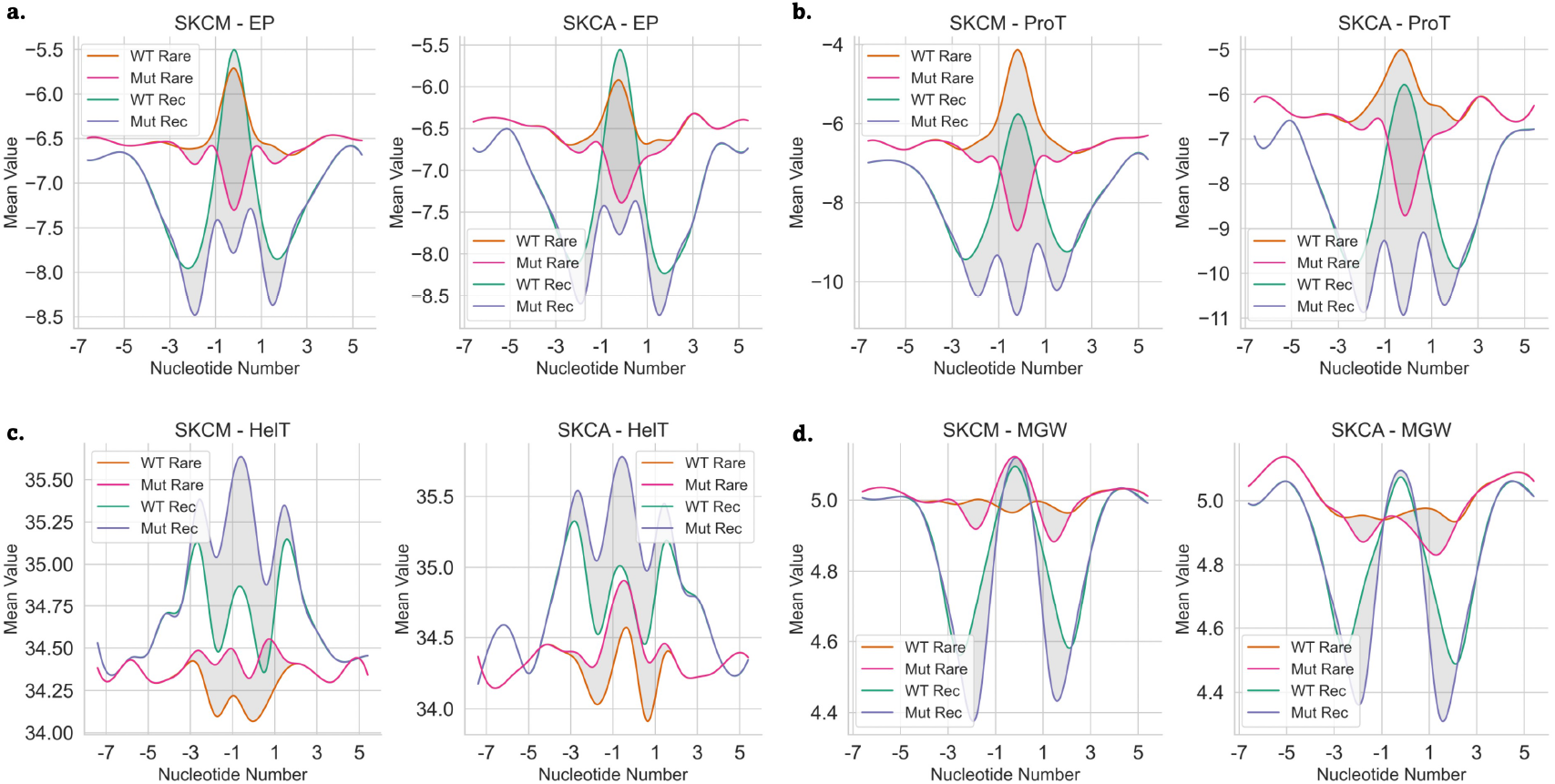
DNA shape results. Using DNAShapR, we predict the **a)** electrostatic potential (EP), **b)** propeller twist (ProT), **c)** helix twist (HT), and **d)** minor groove width (MGW) for nucleotide positions -7 to +7 flanking either side of the single nucleotide variant. Position 0 denotes the location of the substitution. As seen in previous figures, this feature is denoted as “11 feature”. We capture DNA shape values for both the wild-type DNA sequence and the mutant sequence. For both the “rare” (r = 1) and the “recurrent” dataset (r *>* n, where “n” is the threshold value specific to different cancer types), we plot the mean value of all variants for each DNA shape feature and position. The figure presents results for SKCM and SKCA datasets. Recurrent variants tend to occur in regions characterised by high flexibility, bends, and twists, while rare variants are observed in comparatively “flatter” regions. In both classes, there is a shift in DNA properties after the substitution, and this impact regions flanking the substitution site.

#### 3.2.3 Evaluating Top-Performing *CanDrivR-CS* Models on ‘Unseen’ TCGA Data

We evaluated two top-performing models, *CanDrivR-UCEC (Uterine Corpus Endometrial Carcinoma)* and *CanDrivR-SKCM (Skin Cutaneous Melanoma)*, using data from TCGA. Our results showed that *CanDrivR-CS* models generalise well to unseen TCGA datasets (Table 4). *CanDrivR-SKCM* achieved an F1 score of 91.8%. Additionally, *CanDrivR-UCEC* performed well with an F1 score of 79.0% for the TCGA data. Hence, our cancer-specific predictors are also able to generalise to new data with far higher accuracy than our baseline pan-cancer model.

**Table 4:** Performance metrics for *CanDrivR-UCEC* and *CanDrivR-SKCM* models on “seen” (ICGC Test) and “unseen” (TCGA) datasets. Results include accuracy, precision, recall, F1 score, and AUC score.

| Cancer | Dataset | Acc. (%) | Prec. (%) | Rec. (%) | F1 (%) | AUC (%) |
| --- | --- | --- | --- | --- | --- | --- |
| UCEC | ICGC | 86.6 | 87.6 | 85.6 | 86.5 | 93.3 |
| UCEC | TCGA | 81.9 | 82.1 | 76.2 | 79.0 | 82.0 |
| SKCM | ICGC | 87.1 | 84.7 | 89.3 | 86.9 | 87.0 |
| SKCM | TCGA | 87.9 | 93.8 | 90.0 | 91.8 | 87.9 |

## 4 Discussion & Conclusions

In this study, we present *CanDrivR-CS*, a cancer-specific gradient boosting approach designed to distinguish between rare and recurrent missense variants. The primary objective of this work was to investigate whether building cancer-specific machine learning models enhances prediction accuracy, compared to a pan-cancer approach. Our secondary objective was to identify which features are important for distinguishing between rare and recurrent single nucleotide missense variants in cancer genomes. Given the heterogeneity of cancers and their tendency to exhibit a wide variety of rare somatic variants in their genomes, investigating the molecular underpinnings of these variants may shed light on why some variants occur more frequently than others.

Our pan-cancer baseline model demonstrated strong performance, achieving a LOGO-CV F1 score of 79.2%. However, when evaluated on unseen data from the COSMIC database, the model’s F1 score dropped to approximately 65%. This discrepancy can be attributed to the differences in data scale and variant occurrence between the COSMIC and ICGC datasets. The COSMIC dataset is significantly larger than the ICGC dataset, which likely results in certain variants classified as rare (R=1) in the ICGC data appearing more frequently in the COSMIC data. As a consequence, our model, which was trained to identify these variants as rare based on ICGC data, may misclassify them as rare in the COSMIC dataset when they are actually more common (recurrent) there.

In the next part of our analysis, we compared the performance of our pan-cancer baseline model with cancer-specific models. Our findings indicated that developing predictors tailored to specific cancer types significantly improves model performance, compared to models trained on variants across all cancer types. While our baseline model achieved a maximum LOGO-CV F1 score of 79.2%, the *CanDrivR-READ (Rectal Adenocarcinoma)* and *CanDrivR-SKCM* models achieved cross-validation scores of 88% and 90%, respectively, representing an improvement of up to 11%. This improvement underscores the importance of tailoring models to specific cancer types.

While larger datasets generally led to better performance, our results also highlight that factors such as dataset heterogeneity and cancer-specific characteristics may contribute to the overall performance of models. For example, some cancers performed poorly compared to our baseline predictor. Specifically, *CanDrivR-LIAD (Liver Adenocarcinoma)*, and *-PEME (Pediatric Medulloblastoma)* had F1 scores of 56%, and 64%, respectively. This could be attributed to the limited number of variants available for training, as each of these cancer types had fewer than 250 variants. Nevertheless, certain models with a large number of variants still did not perform as well as other cancers. For example, *CanDrivR-MALY (Malignant Lymphoma)* contained 990 variants in total, but only reached an F1 score of 70%. A possible explanation for this might be that malignant lymphoma’s exhibit high genetic heterogeneity (31). MALY has a large number of sub-types including diffuse large B-cell lymphoma (DLBCL), follicular lymphoma, and Hodgkin lymphoma (31) and, each of these subtypes may have unique sets of genetic mutations. Other reasons may be that some lymphomas have a high mutational burdon that could lead to hypermutation, as well as complex genetic landscapes (31; 23). Hence, these complex cancer genomes may be more difficult to model.

In the next part of our work, we investigated feature importance. Interestingly, DNA shape properties such as electrostatic potential, propeller twist, helix twist, and roll consistently ranked among the top features across most cancer types, despite our models being trained on missense variants. Specifically, we found that recurrent variants were typically located in more complex DNA regions, characterised by bends and twists. In contrast, rare variants were more likely to occur in simpler, ‘flatter’ regions. We hypothesise that complex DNA shapes may be more prone to errors during the replication process, potentially acting as mutational hotspots, compared to flatter regions that are easier to replicate. To our knowledge, this is the first study to characterise the prevalence of cancer variants in these types of DNA regions. However, one previous study has shown that G-quadruplex DNA structures are correlated with SCNA breakpoint hotspots (8). We speculate that regions of DNA that are more highly bent or twisted may act as mutational hotspots in both disease and healthy genomes. However, in cancer cells, due to rapid cell division and increased stress on DNA replication and repair mechanisms, these regions may evade detection and repair more easily than in normal tissues.

In this study, we trained models on rare and recurrent cancer variants. Although this approach allowed us to directly compare somatic variants within cancer genomes, it is important to emphasise that due to the definition of our training data, these models are not designed to predict pathogenic and benign variants. The classification of variants as rare or recurrent does not necessarily indicate their role in driving cancer. For instance, recurrent variants that confer a growth advantage may have undergone positive selection across multiple patients due to similar selective pressures (33). Nevertheless, these common variants might only play a role in sustaining cancer cells rather than initiating or driving cancer progression. On the other hand, rare variants could be passenger mutations with no impact on growth or could lead to cancer phenotypes that are not evolutionary stable. This underscores the complex relationship between variant recurrence and their potential roles in cancer.

Therefore, this study primarily serves as a proof of concept, illustrating the advantage of building cancer-specific models and using DNA shape properties when modeling cancer variants. To enhance the interpretability of these findings, exploring alternative methodologies for acquiring high-quality training data to distinguish between pathogenic and neutral variants in a cancer-specific context is essential. Integrating these methods with advanced *in vitro* techniques, such as MAVE, is positioned to advance our understanding of the molecular characteristics of neutral and pathogenic cancer variants and their impact on disease phenotypes.

## 5 Acknowledgments

This work was carried out in the UK Medical Research Council Integrative Epidemiology Unit (MC UU 00032/03) and using the computational facilities of the Advanced Computing Research Centre, University of Bristol. The results published here are based in part on data generated by the TCGA Research Network: https://www.cancer.gov/tcga. For the purpose of open access, the author(s) has applied a Creative Commons Attribution (CC BY) licence to any Author Accepted Manuscript version arising from this submission.

## 6 Funding

This work was funded by Cancer Research UK [C18281/A30905], and supported by resources from the MRC Integrative Epidemiology Unit (MC UU 00032/03).

## Notes

https://github.com/amyfrancis97/CanDrivR-CS

### Competing Interest Statement

The authors have declared no competing interest.

