## Supplementary Material for "*CanDrivR-CS*: A Cancer-Specific Machine Learning Framework for Distinguishing Recurrent and Rare Variants"

### *CanDrivR-CS* Supplementary Material

September 19, 2024

In this Supplementary, we present the details of our optimisation methods for *CanDrivR* and *CanDrivR-CS* models.

#### 1 Software Availability

*CanDrivR-CS* code, training, and test data is available at:
`https://github.com/amyfrancis97/CanDrivR-CS`.

#### 2 Methods & Optimisation

##### 2.1 Optimising the Recurrent Dataset Threshold

###### 2.1.1 *CanDrivR* Baseline Model

Our baseline model classified variants based on whether they are rare or recurrent in the International Cancer Genome Consortium (ICGC) dataset (2). Rare variants were defined as those occurring only once in the ICGC dataset. To optimise the threshold for selecting recurrent variants, we altered our training data to include recurrent variants with donor counts in the range of >1 and >9. Figure 1 shows the mean cross-validation F1 score against the donor count threshold for the recurrent class. For each threshold, we randomly sub-sampled the same number of rare variants ($r = 1$) to create a balanced dataset. We found that a donor count threshold of >2 led to the highest F1 score (approximately 78%), while still maintaining a dataset size of 135,648 missense variants. As a result, we adopted this threshold in the *CanDrivR* baseline model.

###### 2.1.2 *CanDrivR-CS* Models

We then repeated this threshold optimisation process for each of our *CanDrivR-CS* models, adjusting the threshold for each cancer type (Figure 2). For each cancer type, we selected the threshold that resulted in the highest performance.

##### 2.2 Feature Selection

We performed feature selection to identify the most informative features for distinguishing between rare and recurrent mutations. Initially, our baseline XGBoost model used 1,807 features derived from *DrivR-Base* (3). For optimisation purposes, we randomly sampled a balanced training subset of 5,000 variants from the baseline ICGC dataset. The importance of each feature was evaluated using XGBoost's get_booster().get_score function, which ranks features based on their contribution to the model.

In the initial run, 673 out of the 1,807 features were found to be informative; each receiving an importance score of 1 or higher. These 673 features were then ranked by importance, with the top feature being hg48phyloP470way (importance score = 37). To assess how feature inclusion impacted the performance, we employed an iterative approach. Starting with the most important feature, we incrementally added features to the model, performing 673 iterations in total. For each iteration, we plotted the number of features used against the mean F1 score obtained through leave-one-group-out cross-validation (blue line). The standard deviation of the F1 scores across the folds is represented by the orange line. From this, we decided to set a threshold of 351 features for the final model, as including additional features beyond this point did not improve the performance (Figure 3).
From the selected features, we plotted the top 30 based on their importance scores in XGBoost (Figure 4). The most informative features were conservation-based metrics such as PhyloP and PhastCons, alongside DNA-shape properties predicted by DNAShapeR. These DNA-shape properties included Rolls, Helix Twists (HelT), and Minor Groove Width (MGW).

#### 2.3 *CanDrivR-CS*: Comparing F1 Score and Cancer Dataset Size

For each *CanDrivR-CS* model, we derived the F1 cross-validation score. We observed that while some models performed exceptionally well, others did not. To explore this further, we plotted the F1 score against the sample size (Figure 5). A lower-bounding logarithmic curve was then fitted to the lower-bounding points. Although the relationship is not linear, models with poorer performance tended to have smaller training datasets compared to those with better performance. Therefore, we hypothesise that the performance of many of these models would improve with larger datasets available for those particular cancer types.

#### 2.4 *CanDrivR-CS*: How Well Can Cancer-Specific Models Generalise to Other Cancer Datasets?

We aimed to understand the extent to which our cancer-specific models could generalise to other cancer datasets. The rationale behind this testing was for two reasons: firstly, to investigate whether certain cancer types can be pooled into a single predictive model, and secondly, to explore the potential for shared feature group importances across different cancers. By training models on specific cancer types and then testing them on other cancer datasets, we recorded the test accuracy and the F1 score for all models on each dataset. The results for the top models are shown in Figure 6, where the orange point indicates the test results for the cancer dataset used to train the model.
These test results demonstrated that cancer-specific models were able to generalise to other cancer types, but typically with a much lower performance, compared to a cancer-specific context. Across all models, no other datasets showed competing performance with the cancer that they were trained on. This suggests that there is an advantage in building cancer-specific predictors, as each cancer has unique variants that exist within rare and recurrent backgrounds.

#### 2.5 Final Features

In the final section of this supplementary material, we list all of the features used in our final model with a brief description and their sources (see Tables 1 to 6). For a full explanation and description of the feature groups used in *CanDrivR-CS*, please refer to our previous *DrivR-Base* paper and supplementary materials (3).

[3] Amy Francis, Colin Campbell, and Tom R. Gaunt. Drivr-base: a feature extraction toolkit for variant effect prediction model construction. *Bioinformatics*, 40, 3 2024.

[4] Maik Friedel, Swetlana Nikolajewa, Jürgen Sühnel, and Thomas Wilhelm. Diprodb: a database for dinucleotide properties. *Nucleic Acids Research*, 37:D37, 2009.

[5] Luis R. Nassar, Galt P. Barber, Anna Benet-Pagès, Jonathan Casper, Hiram Clawson, Mark Diekhans, Clay Fischer, Jairo Navarro Gonzalez, Angie S. Hinrichs, Brian T. Lee, Christopher M. Lee, Pranav Muthuraman, Beagan Nguy, Tiana Pereira, Parisa Nejad, Gerardo Perez, Brian J. Raney, Daniel Schmelter, Matthew L. Speir, Brittney D. Wick, Ann S. Zweig, David Haussler, Robert M. Kuhn, Maximilian Haeussler, and W. James Kent. The ucsc genome browser database: 2023 update. *Nucleic acids research*, 51:D1188–D1195, 1 2023.

[6] Julien Pelé, Jean Michel Bécu, Hervé Abdi, and Marie Chabbert. Bios2mds: An r package for comparing orthologous protein families by metric multidimensional scaling. *BMC Bioinformatics*, 13:1–7, 6 2012.

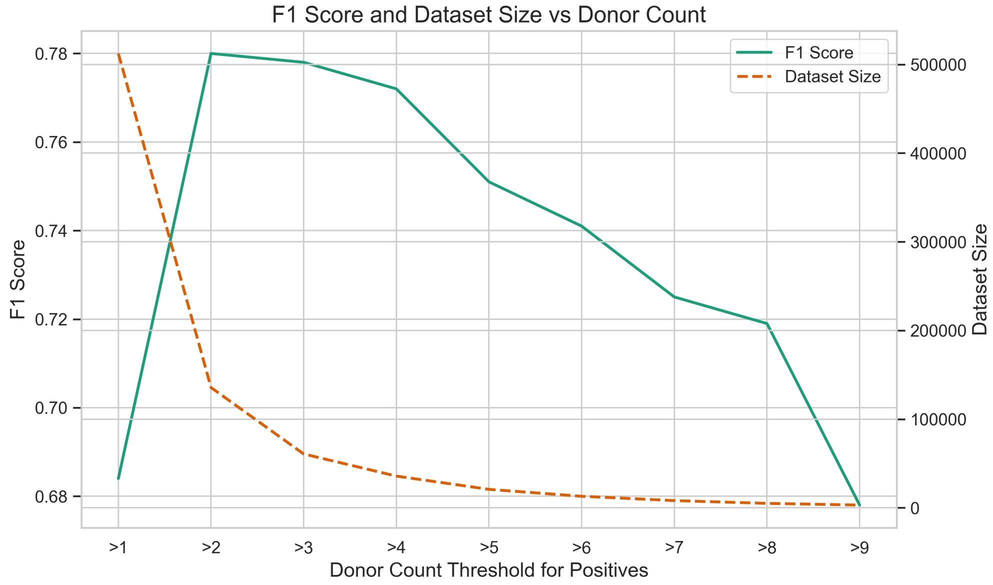


Figure 1: We aimed to establish the best threshold for filtering our recurrent variants. We trained the model using different donor count thresholds ranging from >1 to > 9, and evaluated the models on our cancer-specific test data. We found that filtering variants to those that are found in more than two patients yielded the highest F1 cross-validation score (0.78).

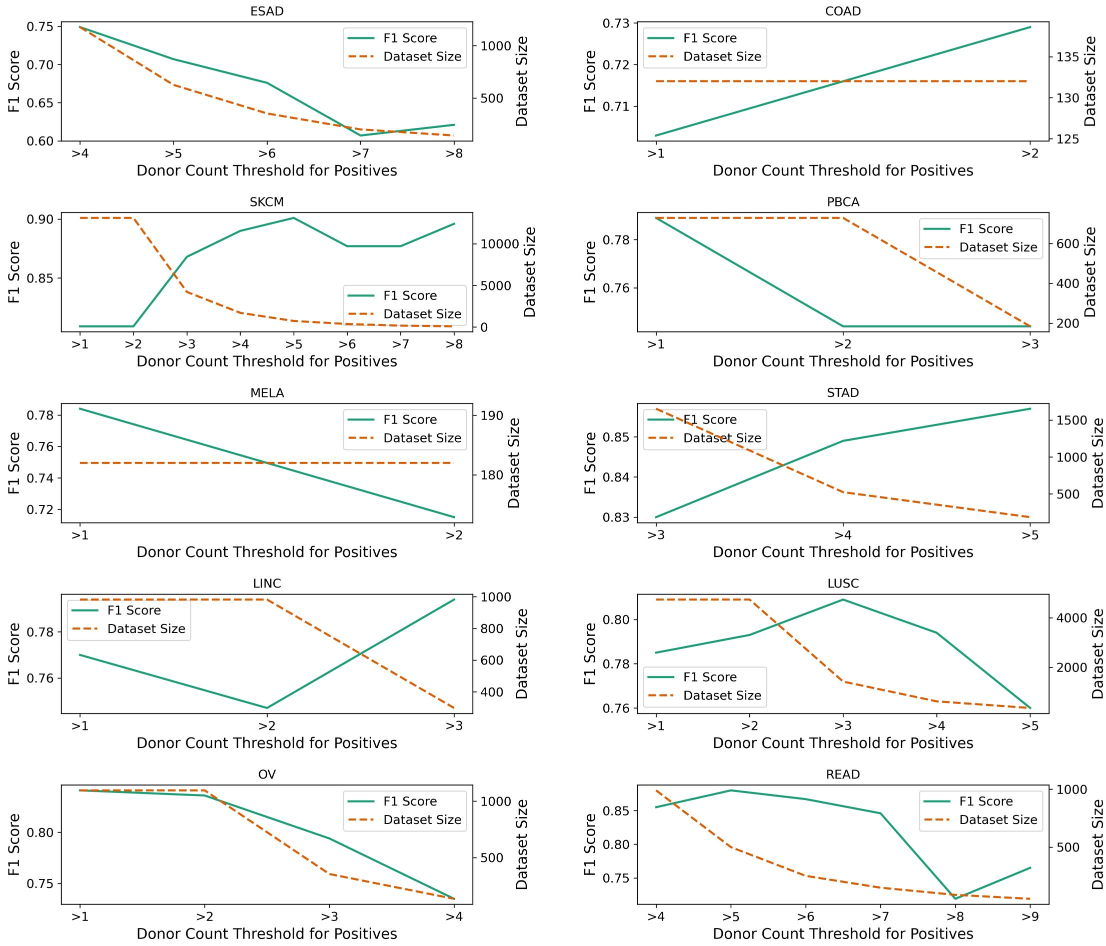


Figure 2: For each cancer type in *CanDrivR-CS*, we optimised the donor count threshold of the recurrent variants. We select the threshold that led to the highest performance for each dataset. Here we show some examples of the optimisation curves produced.

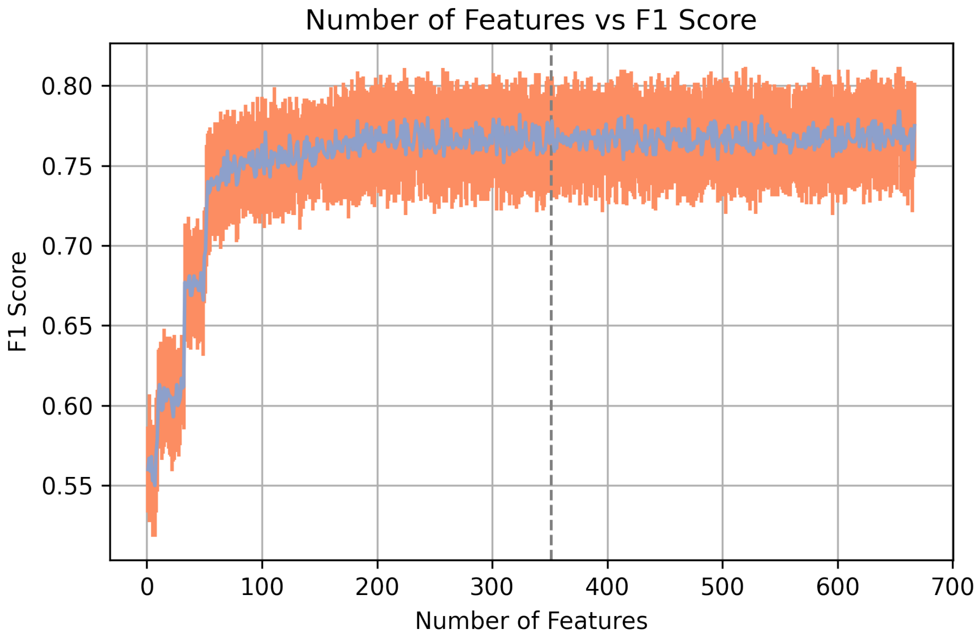


Figure 3: We optimised the features used in our model through sequential feature selection. To reduce the data size for optimisation purposes, we randomly sub-sampled our training dataset to create a balanced subset of 5,000 variants. We then employed XGBoost's get_booster().get_score function to obtain feature importance scores, ranking the top 673 features that had a score greater than 1. Starting with the most important feature, we incrementally added the next most important feature to the model and re-evaluated our baseline model. We plotted the mean cross-validation F1 score for each iteration (in blue) alongside the standard deviation across the folds (in orange). The grey line shows the selected cutoff for diminishing returns.

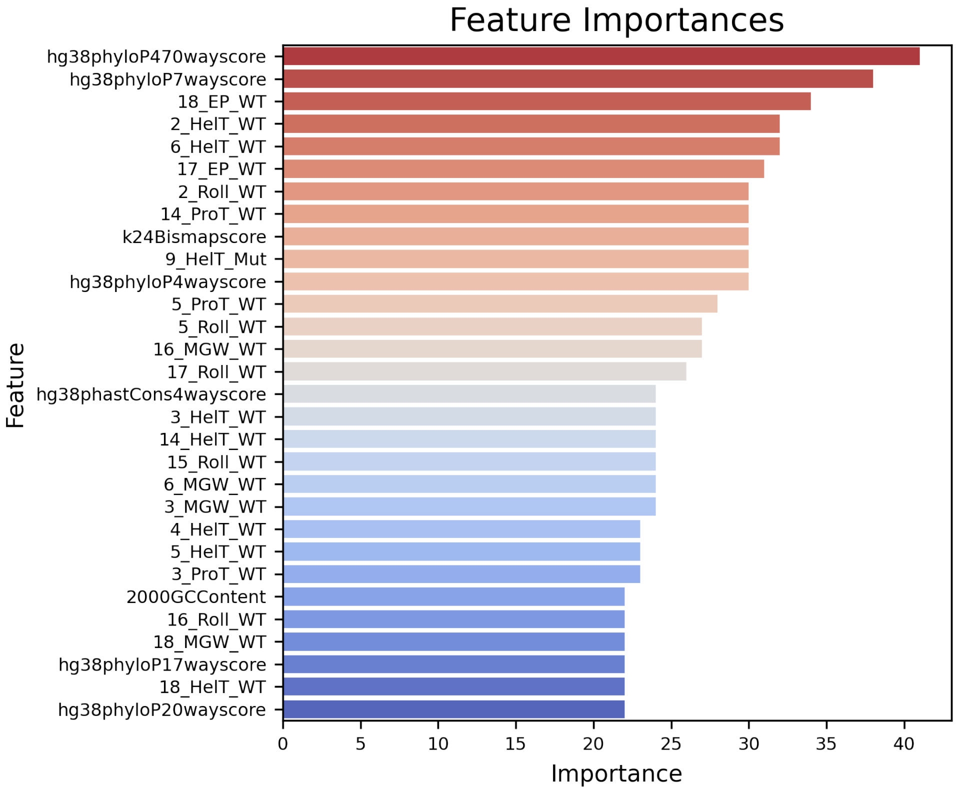


Figure 4: From our features selected in Figure 3, we plotted the top 30 features with their relative importance scores. For more informatio on the feature codes, please refer to supplementary tables 1-6

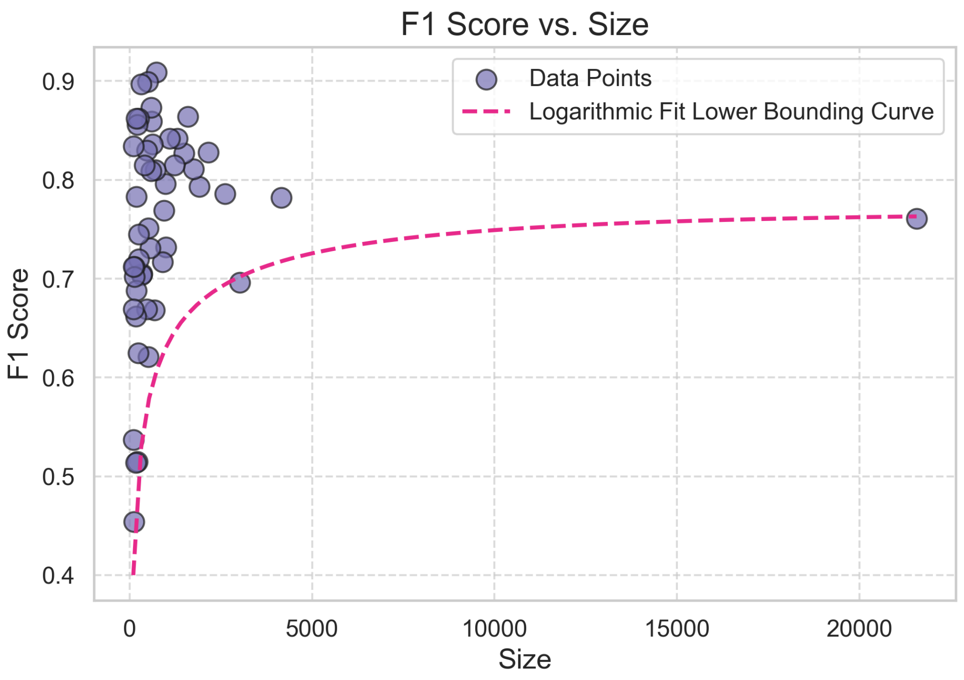


Figure 5: This plot illustrates the relationship between dataset size and F1 score for each of our cancer models in *CanDrivR-CS*. The scatter points represent the F1 scores, while the dashed curve shows the logarithmic fit to the lower bounding F1 scores. There is a non-linear trend, and most poorly-performing models had smaller training datasets.

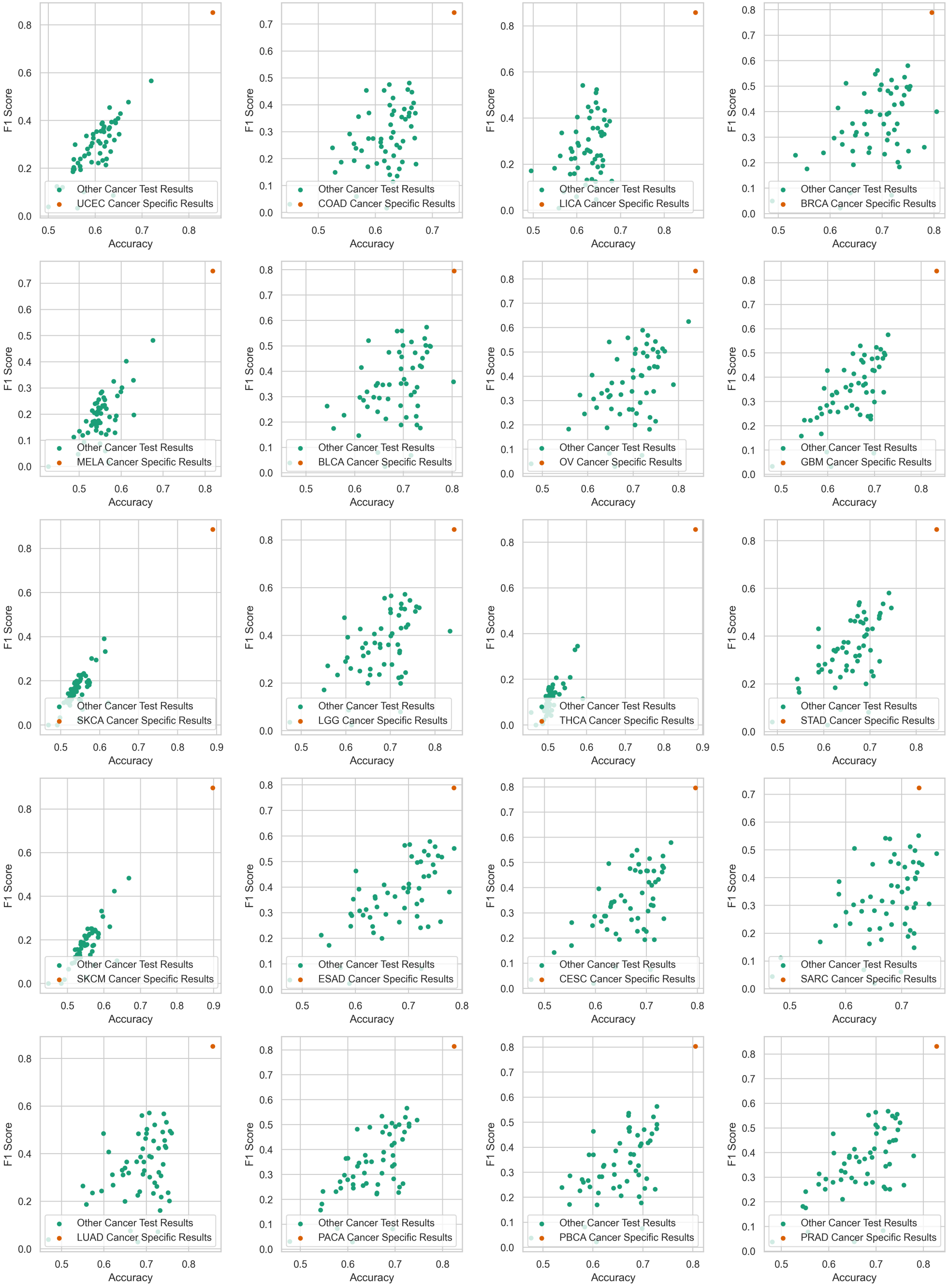


Figure 6: Here, we plotted the cancer-specific cross-validation F1 score vs accuracy against the results of testing on all other cancer datasets. Specifically, we trained our *CanDrivR-CS* models, and tested each model on all other cancer datasets. The reasoning for this was to test the ability of our models to generalise to other cancer datasets. We found that for each cancer model, the corresponding cancer test dataset gives the best performance. No other cancer datasets are competitive or even close to the performance of the cancer data that was used to train the model. Hence, we cannot pool any cancer-specific models together.

| Code | Positions | Description |
|---|---|---|
| **Wild Type (WT)** | | |
| HelT_WT | 2-18 | Helical Twist |
| Roll_WT | 2-18 | Roll |
| ProT_WT | 2-18 | Propeller Twist |
| EP_WT | 2-18 | Electrostatic Potential |
| MGW_WT | 2-8 | Minor Groove Width |
| **Mutant (Mut)** | | |
| HelT_Mut | 9,-13 | Helical Twist |
| Roll_Mut | 8-13 | Roll |
| ProT_Mut | 9-13 | Propeller Twist |
| EP_Mut | 9-13 | Electrostatic Potential |
| MGW_Mut | 9-13 | Minor Groove Width |

Table 1: This table shows the DNA shape feature used in our final model. They include helix twists, propeller twists, electrostatic potential, and minor groove width. All of our DNA shape features were sourced from the R package 'DNAShapeR' (1).

| Feature Code | Description |
|---|---|
| **PhyloP Scores** | |
| hg38phyloP470wayscore | PhyloP score across 470 vertebrate species |
| hg38phyloP17wayscore | PhyloP score across 17 vertebrate species |
| hg38phyloP7wayscore | PhyloP score across 7 vertebrate species |
| hg38phyloP100wayscore | PhyloP score across 100 vertebrate species |
| hg38phyloP4wayscore | PhyloP score across 4 vertebrate species |
| hg38phyloP30wayscore | PhyloP score across 30 vertebrate species |
| hg38phyloP20wayscore | PhyloP score across 20 vertebrate species |
| **PhastCons Scores** | |
| hg38phastCons4wayscore | PhastCons score across 4 vertebrate species |
| hg38phastCons17wayscore | PhastCons score across 17 vertebrate species |
| hg38phastCons30wayscore | PhastCons score across 30 vertebrate species |
| hg38phastCons20wayscore | PhastCons score across 20 vertebrate species |
| hg38phastCons470wayscore | PhastCons score across 470 vertebrate species |
| hg38phastCons7wayscore | PhastCons score across 7 vertebrate species |
| hg38phastCons100wayscore | PhastCons score across 100 vertebrate species |
| **Mappability Scores** | |
| k24Bismapscore | Bismap score (24-mer) |
| k36Bismapscore | Bismap score (36-mer) |
| k100Bismapscore | Bismap score (100-mer) |
| k100Umapscore | Umap score (100-mer) |
| k50Umapscore | Umap score (50-mer) |

Table 2: These features represent the conservation-based and mappability scores used in our final models. We downloaded conservation measures from the UCSC Genome Browser (5).

| Feature Code | Description |
|---|---|
| **GC Content** | |
| $< n >$GCContent | where 'n' is the GC content calculated over 20, 40, 60, 80, 100, 200, 500, 1000, 2000 base pair windows |
| **CpG Count** | |
| $< n >$CpGCount | where 'n' is the CpG count calculated over 20, 60, 100, 200, 500, 1000, 2000 base pair windows |
| **Observed/Expected CpG Ratio (CpG_obs_exp)** | |
| $< n >$CpG_obs_exp | Observed/expected CpG ratio, where 'n' is 20, 40, 60, 80, 100, 200, 500, 1000, 2000 base pair windows |

Table 3: This table presents the GC content, CpG count, and observed vs expected CpG ratio for window sizes ranging between 20-2000 base pairs. For a full description of these features and how they were calculated, please refer to our *DrivR-Base* paper (3).

| Feature Code | Description |
|---|---|
| 10_3_x | Spectrum feature for a window size of 10 and k-mer size of 3 |
| 10_2_x | Spectrum feature for a window size of 10 and k-mer size of 2 |
| 4_2_x | Spectrum feature for a window size of 4 and k-mer size of 2 |
| 10_1_x | Spectrum feature for a window size of 10 and k-mer size of 1 |
| 8_2_x | Spectrum feature for a window size of 8 and k-mer size of 2 |
| 6_1_x | Spectrum feature for a window size of 6 and k-mer size of 1 |
| 8_1_x | Spectrum feature for a window size of 8 and k-mer size of 1 |
| 2_1_w | Spectrum feature for a window size of 2 and k-mer size of 1 |
| 10_2_z | Spectrum feature for a window size of 10 and k-mer size of 2 |
| 8_1_z | Spectrum feature for a window size of 8 and k-mer size of 1 |
| 10_1_z | Spectrum feature for a window size of 10 and k-mer size of 1 |
| 6_1_z | Spectrum feature for a window size of 6 and k-mer size of 1 |
| 8_2_z | Spectrum feature for a window size of 8 and k-mer size of 2 |
| 6_2_z | Spectrum feature for a window size of 6 and k-mer size of 2 |
| 4_1_z | Spectrum feature for a window size of 4 and k-mer size of 1 |
| 10_3_z | Spectrum feature for a window size of 10 and k-mer size of 3 |
| 6_3_z | Spectrum feature for a window size of 6 and k-mer size of 3 |
| 6_5_z | Spectrum feature for a window size of 6 and k-mer size of 5 |

Table 4: Feature codes and descriptions for kernel-based spectrum features. The features were derived using k-mer and window size combinations from wild type and mutant sequences. Please see our *DrivR-Base* paper for a detailed description of the calculations (3)

| Feature Code | Description |
|---|---|
| JTT | Jones-Taylor-Thornton (JTT) substitution matrix |
| PAM40 | Point Accepted Mutation (PAM) matrix, 40% divergence |
| GONNET | Gonnet substitution matrix |
| BLOSUM30 | BLOSUM (BLOcks Substitution Matrix) for sequences with 30% identity |
| BLOSUM80 | BLOSUM (BLOcks Substitution Matrix) for sequences with 80% identity |
| BLOSUM45 | BLOSUM (BLOcks Substitution Matrix) for sequences with 45% identity |
| JTT_TM | Jones-Taylor-Thornton (JTT) matrix for transmembrane proteins |
| PHAT | PHAT (Percent Hit Acceptance Threshold) substitution matrix |

Table 5: Feature codes and descriptions for substitution matrices. We show the substitution matrices that were used as features in our final model, all of which where taken from the Bio2mds package in R (6).

| **Feature Code** | **Description** |
|---|---|
| Inclination | Positions: left and right of the variant (WT and Mutant) |
| Direction | Positions: left and right of the variant (WT and Mutant) |
| Probability_contacting_nucleosome_core | Positions: left and right of the variant (WT) |
| Stacking_energy | Positions: left and right of the variant (WT, RNA) |
| Twist_tilt | Positions: left and right of the variant (WT) |
| Twist_shift | Positions: left and right of the variant (WT and Mutant) |
| Tilt_roll | Positions: left and right of the variant (Mutant) |
| Flexibility_shift | Positions: left and right of the variant (WT) |
| Hydrophilicity_(RNA) | Positions: left and right of the variant (Mutant) |
| Minor_Groove_Size | Positions: left and right of the variant (WT and Mutant) |
| Entropy_(RNA) | Positions: left and right of the variant (WT) |
| Tilt_(RNA) | Positions: left and right of the variant (WT and Mutant) |
| Bend | Positions: left and right of the variant (WT and Mutant) |
| Shift_shift | Positions: right of the variant (WT) |
| Twist_twist | Positions: right of the variant (Mutant) |
| Shift_slide | Positions: right of the variant (Mutant) |
| Minor_Groove_Depth | Positions: right of the variant (Mutant) |
| Propeller_Twist | Positions: left of the variant (Mutant) |
| Tilt_(DNA-protein_complex) | Positions: left of the variant (Mutant) |
| Tip | Positions: left of the variant (Mutant) |
| Guanine_content | Positions: right of the variant (WT) |
| Thymine_content | Positions: right of the variant (WT) |
| Major_Groove_Distance | Positions: right of the variant (WT) |
| Slide_rise | Positions: right of the variant (WT) |
| Wedge | Positions: left of the variant (WT) |
| Shift_(DNA-protein_complex) | Positions: left of the variant (WT) |
| Slide_(RNA) | Positions: left of the variant (WT) |
| Tilt_roll | Positions: left of the variant (WT) |
| Twist_slide | Positions: left of the variant (WT) |
| Major_Groove_Width | Positions: left of the variant (WT) |

Table 6: Feature codes and descriptions for dinucleotide properties from DiProDB (4). For a detailed description on the position of the feature, please refer to *DrivR-Base* supplementary material (3).
